# Genome assemblies and annotations of the poneroid ants *Platythyrea mocquerysi* (Ponerinae) and *Stigmatomma oregonense* (Amblyoponinae)

**DOI:** 10.64898/2026.10.08.757030

**Authors:** Adria C. LeBoeuf, Joel Vizueta, Zijun Xiong, Yuqi Reitsema-Wang, Andrew Brown, Jean-Jacques Rafanomezantsoa, Helder Hugo, Brian L. Fisher

## Abstract

We present draft genome assemblies for two poneroid ant species with distinctive life history traits: *Platythyrea mocquerysi* Emery, 1899 (Ponerinae) and *Stigmatomma oregonense* Wheeler, 1915 (Amblyoponinae). *Platythyrea mocquerysi* is a Malagasy ant in which larvae produce secretions from a ventral tubercle that are consumed by adults, representing an unusual form of larva-to-adult trophallaxis. *Stigmatomma oregonense*, a North American “dracula ant,” feeds non-lethally on larval haemolymph. Using PacBio HiFi long-read sequencing, we assembled the *P. mocquerysi* genome (187 Mb, 307 scaffolds, N50 = 5.26 Mb, BUSCO 99.2% complete) and the *S. oregonense* genome (262 Mb, 1,189 scaffolds, N50 = 0.84 Mb, BUSCO 98.3% complete). These genomes encode 10,860 and 12,354 protein-coding genes, respectively. Repeat content is higher in S. oregonense than in P. mocquerysi, 54.3% against a lower bound of 15.7%. Most of the difference is in unclassified repeats, which we show to be diverged transposable elements. Comparative analysis with two additional poneroid genomes from closely related species reveals that genome size variation across poneroid ants is largely explained by transposable element dynamics, with notable expansions in several gene families including carboxylesterases, serine proteases, and lipid metabolism genes in Amblyoponinae. These genomes provide essential resources for investigating the molecular basis of unconventional brood management strategies in ants.

## Introduction

Poneroid ants display a remarkable range of life history traits, with *Platythyrea mocquerysi* Emery, 1899 (Ponerinae) and *Stigmatomma oregonense* Wheeler, 1915 (Amblyoponinae) serving as particularly distinctive examples (Emery, 1899; Brown, 1975; Wheeler, 1915; Bolton, 1995; Yoshimura & Fisher, 2012). In *P. mocquerysi*, larvae develop a specialised ventral tubercle that secretes fluids consumed by adults, a rare phenomenon among ants that may influence colony nutrition and social dynamics (Villet et al., 1990). *Stigmatomma oregonense*, belonging to the group known as “dracula ants,” is noted for a unique behaviour in which workers feed non-lethally on larval haemolymph by making small punctures, a strategy seldom observed in other ant groups (Brown, 1960, 1975; Yoshimura & Fisher, 2012). Both species also exhibit cryptic nesting habits; *P. mocquerysi* is typically found in decaying wood, while *S. oregonense* occupies both decaying wood and subterranean environments (Ward, 1988).

The genetic mechanisms underlying these ants’ adaptations remain largely unknown. Previous genomic studies have focused mainly on more conventional ant species, leaving these unusual lineages understudied. The Global Ant Genomics Alliance (GAGA; Vizueta et al., 2025) and others (e.g. Romiguier et al., 2022) have built a powerful backbone for genomic studies in ants, but detailed genome reports for individual species with distinct life history and social traits remain essential for enabling targeted studies of gene function and evolution.

Here we present high-quality genome assemblies and annotations for *P. mocquerysi* and *S. oregonense* (Figure 1), generated using PacBio HiFi long-read sequencing and annotation following the GAGA consortium pipeline. We place these genomes in a comparative context with two additional poneroid species from the GAGA repository, *Platythyrea punctata* (mostly clonal) and *Stigmatomma* sp. (an undescribed species from China), to characterise genome size variation, repeat landscape dynamics, and gene family evolution across these species in the poneroid clade.

**Figure 1.**
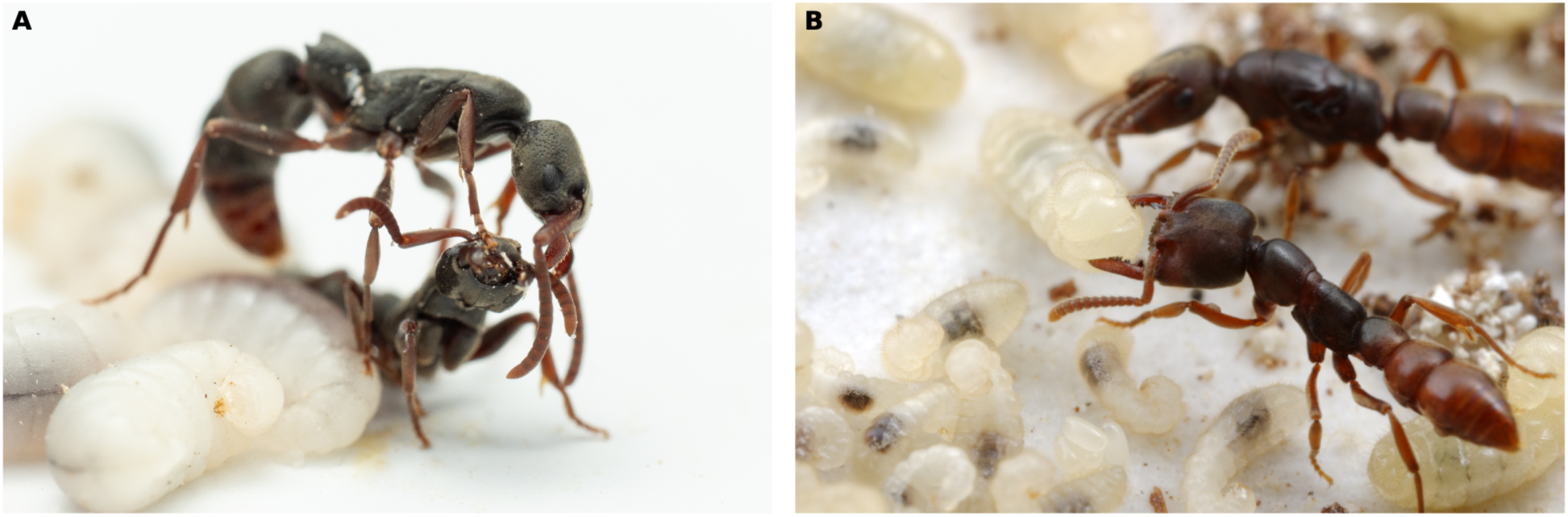
The two poneroid ant species sequenced in this study. (A) *Platythyrea mocquerysi* (Ponerinae), dominance battle over brood. (B) *Stigmatomma oregonense* (Amblyoponinae), queen and worker with brood. Photographs by François Brassard (A) and Alexander Wild (B), reproduced with permission.

## Methods & Materials

### Statement of Legal Access and Benefit-Sharing (ABS) Compliance

*Platythyrea mocquerysi* specimens were collected in Madagascar under research permit N° 073/23/MEDD/SG/DGGE/DAPNRE/SCBE.Re (10 March 2023, in the name of Andrew Francis Brown) and exported under permit N° 187N-EA05/MG23 (03 May 2023). *Stigmatomma oregonense* was collected in Canada, where no Nagoya Protocol requirements apply.

### Sample Collection

*Platythyrea mocquerysi* were collected from Madagascar: Toliara Province, Menabe Region: Forêt de Kirindy, 20.07216°S, 44.67268°E, 38 m elevation, 21 April 2023, tropical dry forest (research permit N° 073/23/MEDD/SG/DGGE/DAPNRE/SCBE.Re).

*Stigmatomma oregonense* colonies were collected from Canada: British Columbia, Vancouver Island, Heart Lake, 48.973781°N, 123.826187°W, October 2023, from semi-rotten logs. Colonies were maintained at the Social Fluids Lab at the University of Fribourg at 25 C and 60% humidity until use.

### DNA Extraction

For *P. mocquerysi*, two live pupae were collected from colony

23Pmoc24/BLF42828/CASENT0872487. Cocoons were carefully removed before pupae were prepared for DNA extraction. For *S. oregonense*, three late-stage larvae from colony 23Sore.44/CASENT0872488 had their gut carefully removed before DNA extraction. High molecular weight (HMW) genomic DNA was extracted using a Qiagen Gentra Puregene Tissue Kit (Qiagen, cat. no. 158622) following the manufacturer’s guidelines for tissue samples (Qiagen Gentra Puregene Handbook 06/2011). Samples were homogenised using a microcentrifuge tube pestle. The 10x Genomics sample preparation protocol for DNA extraction from single insects was employed. Prior to library preparation, genomic DNA was assessed for quantity using a Qubit 4.0 fluorometer (Qubit dsDNA BR Assay kit, Q32850, Thermo Fisher Scientific), for quality using an Advanced Analytical FEMTO Pulse instrument (Genomic DNA 165 kb Kit, FP-1002-0275, Agilent), and for purity using a DeNovix DS-11 UV-Vis spectrophotometer.

### Library Preparation

SMRTbell libraries were prepared according to the PacBio guideline “Preparing whole genome and metagenome libraries using SMRTbell® prep kit 3.0” (Part Number 102-166-600, March 2023). Briefly, HMW gDNA was sheared using a Megaruptor 3, then concentrated and cleaned using 1× SMRTbell clean-up beads. Sheared DNA was assessed for quantity using a Qubit 4.0 fluorometer (Qubit dsDNA HS Assay kit, Q32851) and for quality/size distribution using a FEMTO Pulse instrument (target range 13–17 kb). The standard SMRTbell protocol was followed, including end-repair and A-tailing, ligation of barcoded overhang adapters, purification with AMPure PB beads, and nuclease treatment. Final libraries were checked for quantity and quality using a Qubit 4.0 fluorometer and FEMTO Pulse instrument.

For *P. mocquerysi*, the standard SMRTbell prep kit 3.0 protocol was used with sufficient input DNA. For *S. oregonense*, the Ultra-Low DNA Input protocol was employed due to limited material availability (approximately 20 ng input DNA).

### Sequencing

For *P. mocquerysi* (PB131), two pooled barcoded libraries were sequenced on a PacBio Sequel IIe instrument in September 2023 at the Next Generation Sequencing Platform, University of Bern. For *S. oregonense* (PB140), three pooled libraries were sequenced on a PacBio Revio instrument in November 2023 at the same facility. Both runs generated HiFi (circular consensus sequencing) reads.

### RNA Sequencing

RNA-seq data were generated to assist genome annotation. For *P. mocquerysi*, RNA was extracted from three sample types from colony 23Pmoc29: pupae (n = 2), adult males (n = 2), and late-instar larvae (n = 4). For *S. oregonense*, RNA was extracted from two sample types from colony Sore.44: adult workers (n = 3) and late-instar larvae (n = 3). RNA-seq libraries were prepared and sequenced by BGI.

### Genome Assembly

PacBio HiFi reads were assembled using both Wtdbg2 v2.5 (parameters: “-t 16 -x ccs -g 300m”; Ruan and Li, 2020) and Hifiasm v0.19.8-r603 (default parameters; Cheng et al., 2021). Assemblies were evaluated based on contiguity metrics and gene completeness with Compleasm v0.2.6 (Huang and Li, 2023) using the BUSCO Hymenoptera_odb12 dataset (n = 5,920). The Hifiasm assembly was selected for *S. oregonense*, while the Wtdbg2 assembly following the GAGA assembly pipeline (Vizueta et al., 2025) was chosen for *P. mocquerysi*. Duplicated scaffolds were identified and filtered using the Funannotate “clean” pipeline v1.8.3 (https://github.com/nextgenusfs/funannotate). Assemblies were screened for putative contamination using the NCBI Foreign Contamination Screen tool FCS-GX v0.5 (Astashyn et al., 2024).

Both species were sequenced with HiFi chemistry and assembled with both assemblers. For *P. mocquerysi* the Wtdbg2 assembly was far more contiguous (N50 5.26 Mb against below 1 Mb) and was retained, following the GAGA pipeline in which contigs are scaffolded with SSPACE-LongRead and polished with NextPolish. For *S. oregonense* the Wtdbg2 assembly was substantially smaller than expected for the genome, and the primary Hifiasm haplotype was retained without further scaffolding or correction. The Hifiasm assemblies of *P. mocquerysi*, which span 297.0 and 254.2 Mb, are reported in Table S1.

### Genome Annotation

Genome annotation was conducted following the pipeline optimised for ant genomes generated by the GAGA project (Vizueta et al., 2025). Repeats were identified and masked using RepeatMasker v4.1.4 (Smit et al., 2013–2015) based on known and *de novo* repeat libraries.

General gene annotations were conducted by combining gene models from several sources. First, RNA-seq data were aligned to the reference repeat soft-masked genome assembly using STAR v2.7.2b (Dobin et al., 2013) with default parameters. Publicly available gene annotations from *Drosophila melanogaster*, *Tribolium castaneum*, *Nasonia vitripennis*, *Apis mellifera*, *Ooceraea biroi*, and *Camponotus floridanus* were used to conduct homology-based gene predictions using GeMoMa v1.7.1 (Keilwagen et al., 2016, 2019), which also incorporated the RNA-seq evidence for splice site prediction. Second, independent RNA-seq alignments were merged to create a consensus GTF using StringTie v2.1.5 (Kovaka et al., 2019), after which BestORF (Molquest package, Softberry) was used to identify open reading frames (ORFs) in the transcript sequences; transcripts with incomplete ORFs were filtered out. Third, approximately 1,000 high-quality genes from the GeMoMa prediction were used to train Augustus v3.2.2 (Stanke et al., 2006). *De novo* gene prediction was then performed using Augustus with the repeat-masked genome, filtering out genes shorter than 150 bp or with incomplete ORFs. Finally, gene annotations from the three sources of evidence were combined by merging them into a single GFF using an in-house Perl script, and transposon-related proteins were identified and filtered using BLASTP v2.2.31 (Camacho et al., 2009) against the SwissProt database and the transposable element protein database from RepeatMasker.

Specific gene families of interest in ant biology were identified and reannotated using BITACORA v1.3 (Vizueta et al., 2020), except for odorant receptors (ORs) and gustatory receptors (GRs), for which HAPpy-ABCENTH v1.0 (https://github.com/biorover/HAPpy-ABCENTH) was used. ORs and GRs are reported as intact when the model lacks the fragmentary marker used by HAPpy-ABCENTH, a class that includes a minority scored Partial or Pseudogene. That marker was not applied in the *P. punctata* and *Stigmatomma* sp. runs, so fragment counts are comparable only between *P. mocquerysi* and *S. oregonense*.

Gene model completeness was also assessed at the protein level with Compleasm v0.2.6 in protein mode, using the representative protein set for each species. Values are reported against hymenoptera_odb12 dataset (n = 5,920); the same analysis against hymenoptera_odb10 dataset (n = 5,991) gave concordant values.

### Comparative Genomic Analysis

To contextualise the two new genome assemblies, we compared them with the two available poneroid species from the same genus generated by GAGA (Vizueta et al., 2025): *Platythyrea punctata* (GAGA-0090; Ponerinae) and *Stigmatomma* sp. (GAGA-0391; Amblyoponinae). Assembly statistics were computed from FASTA files. Repeat content was quantified from RepeatMasker GFF annotation files, with repeat elements classified into broad categories (LTR elements, DNA transposons, LINEs, SINEs, rolling circles, simple/tandem repeats, and unclassified) based on the Class= attribute. Gene family counts were extracted from the full GFF3 annotation files: OR and GR counts from HAPpy-ABCENTH annotations, and IR, OBP, CSP, and other families from BITACORA annotations.

Given the more fragmented assembly of *S. oregonense*, its expanded gene families were tested for assembly artefacts in four ways. First, gene coordinates were taken from the annotation GFF3 and scaffold lengths from the assembly FASTA, and the proportion of genes lying within 5 kb of either scaffold terminus was compared with the null expectation (n_scaffolds × 2 × 5,000) / genome size, which is 4.5% for this assembly. Second, coding sequence was reconstructed for every carboxylesterase, acyl-CoA desaturase and odorant receptor model by concatenating its CDS features in coordinate order and reverse-complementing minus-strand models, then checked for an ATG start, a canonical stop codon (TAA, TAG or TGA) and a length divisible by three; odorant receptors were partitioned by the HAPpy-ABCENTH Status attribute. Third, predicted protein lengths were derived from summed CDS length for odorant receptors, whose complete forms are length-constrained. Fourth, family sizes were compared with those annotated in the independently assembled *Stigmatomma* sp. genome (GAGA-0391).

Genome size was also estimated from the raw HiFi reads without reference to an assembly. Canonical k-mer frequencies in the reads were fitted with GenomeScope 2.0 (Ranallo-Benavidez et al., 2020), and a complementary model-free estimate was computed as total k-mer abundance divided by the modal genomic k-mer coverage.

Repeat collapse in the *P. mocquerysi* assembly was quantified by comparing canonical k-mer copy number (k = 21) between that assembly and each Hifiasm haplotype of the same reads, on a 1/64 hash subsample so that multiplicities scale to unbiased genome-wide totals. Sequence was partitioned by copy number in the Hifiasm assembly and the proportion retained by the Wtdbg2 assembly computed for each class. Both Hifiasm assemblies span more sequence than the deposited one (297.0 and 254.2 Mb) and bracket the GenomeScope 2 estimate. Against either haplotype, unique sequence is 96% retained by the Wtdbg2 assembly and sequence at 20 to 99 copies only 7 to 9% retained, and the longest low-complexity array is 52 kb in the deposited assembly against 214 and 260 kb in the Hifiasm assemblies (Table S1).

## Results and Discussion

### Genome Assembly Quality

The *P. mocquerysi* genome assembled into 307 scaffolds totalling 187.1 Mb, with an N50 of 5.26 Mb (L50 = 15) and a GC content of 48.0% (Table 1; Figure 2). The *S. oregonense* genome assembled into 1,189 scaffolds totalling 261.3 Mb, with an N50 of 0.84 Mb (L50 = 77) and a GC content of 44.8%. BUSCO completeness assessment using Compleasm with the Hymenoptera_odb12 dataset (n = 5,920) showed 99.2% complete genes (98.9% single-copy, 0.4% duplicated) for *P. mocquerysi* and 98.3% complete (96.2% single-copy, 2.1% duplicated) for *S. oregonense*. The lower contiguity of the *S. oregonense* assembly reflects the ultra-low DNA input (approximately 20 ng) used for library preparation, which limited coverage depth. NCBI FCS-GX contamination screening identified 16 scaffolds in *S. oregonense* (15 bacterial [Firmicutes] and 1 mitochondrial) that were excluded from the final assembly.

**Figure 2.**
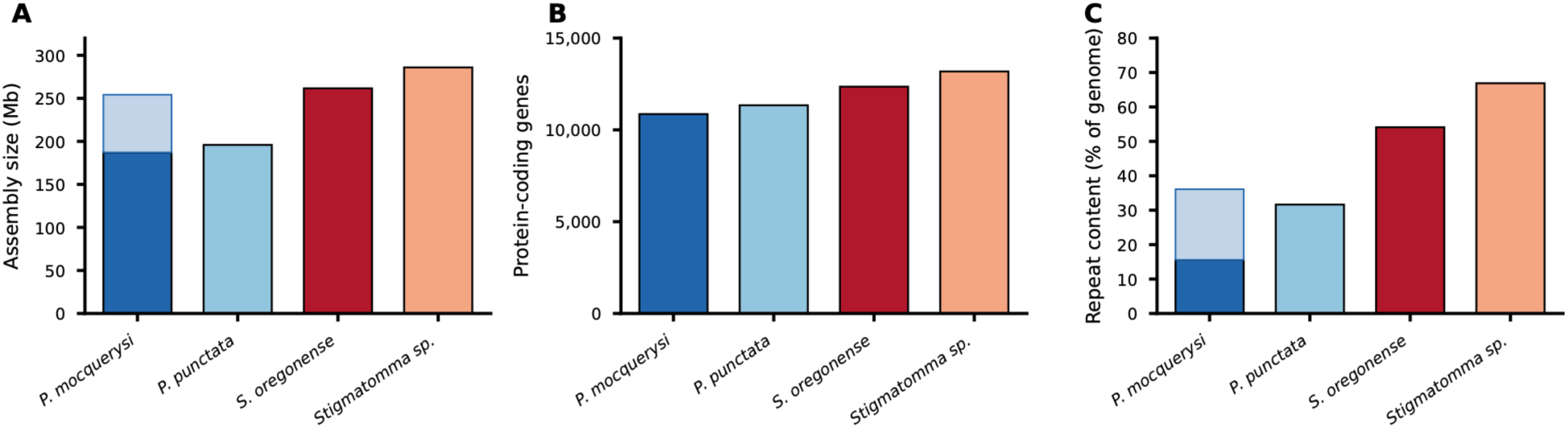
Genome overview of four poneroid ant species. (A) Genome size in megabases. (B) Number of representative protein-coding genes. (C) Total repeat content as percentage of genome. Blue shades indicate Ponerinae; red shades indicate Amblyoponinae. In (A) and (C) the lighter upper portion of the *P. mocquerysi* bar is sequence inferred from k-mer analysis of the reads but not recovered by the assembly.

**Table 1.**
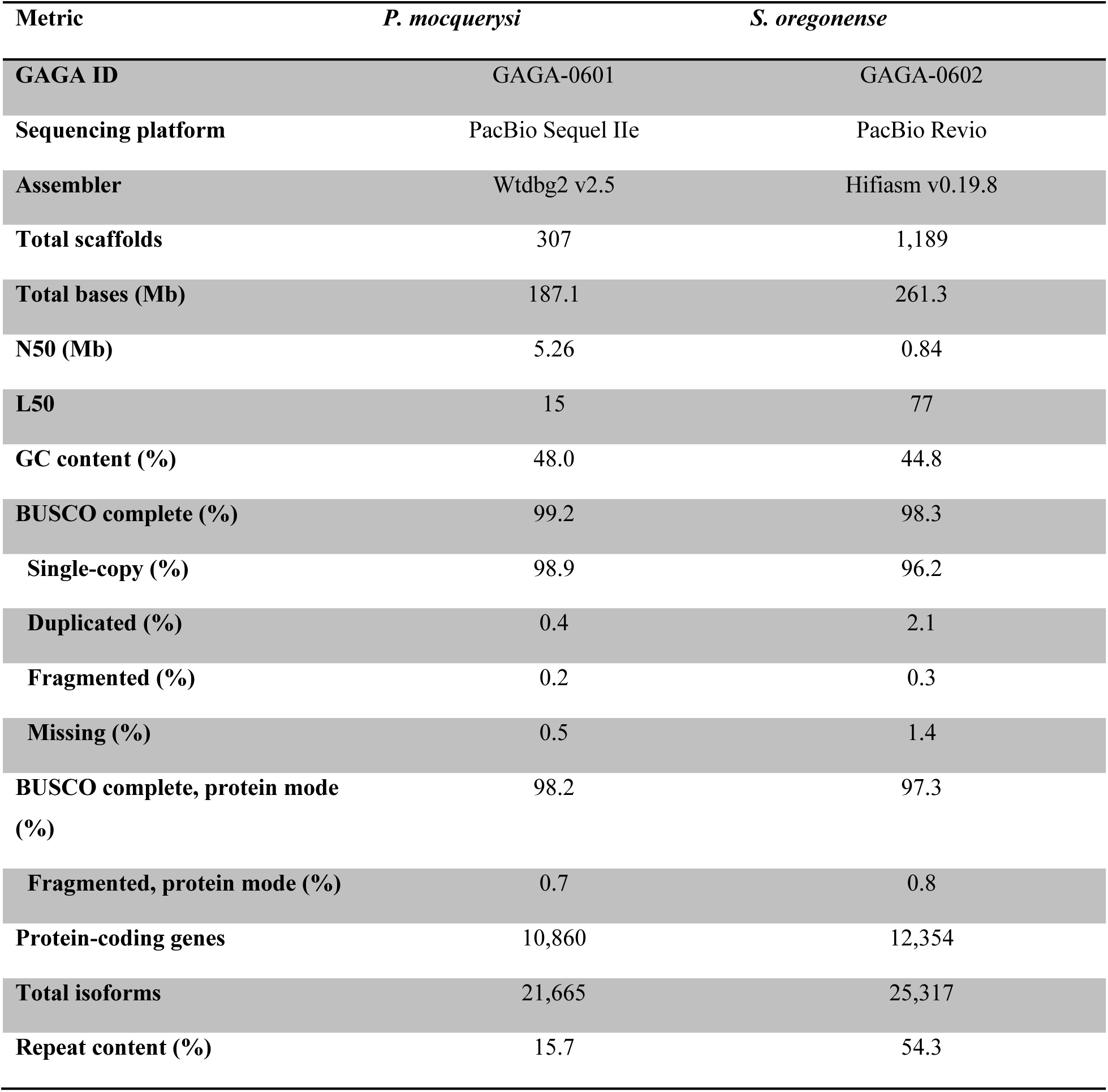
Genome assembly and annotation statistics for *P. mocquerysi* and *S. oregonense*. BUSCO completeness was assessed with Compleasm using the Hymenoptera_odb12 dataset (n = 5,920).

| Metric | *P. mocquerysi* | *S. oregonense* |
| --- | --- | --- |
| **GAGA ID** | GAGA-0601 | GAGA-0602 |
| **Sequencing platform** | PacBio Sequel IIe | PacBio Revio |
| **Assembler** | Wtdbg2 v2.5 | Hifiasm v0.19.8 |
| **Total scaffolds** | 307 | 1,189 |
| **Total bases (Mb)** | 187.1 | 261.3 |
| **N50 (Mb)** | 5.26 | 0.84 |
| **L50** | 15 | 77 |
| **GC content (%)** | 48.0 | 44.8 |
| **BUSCO complete (%)** | 99.2 | 98.3 |
| **Single-copy (%)** | 98.9 | 96.2 |
| **Duplicated (%)** | 0.4 | 2.1 |
| **Fragmented (%)** | 0.2 | 0.3 |
| **Missing (%)** | 0.5 | 1.4 |
| **BUSCO complete, protein mode (%)** | 98.2 | 97.3 |
| **Fragmented, protein mode (%)** | 0.7 | 0.8 |
| **Protein-coding genes** | 10,860 | 12,354 |
| **Total isoforms** | 21,665 | 25,317 |
| **Repeat content (%)** | 15.7 | 54.3 |

Assembly-free estimates agreed with the assembly for *S. oregonense* but not for *P. mocquerysi*. For *S. oregonense*, a model-free k-mer estimate of 263 Mb lies within 1% of the 261.3 Mb assembly, and GenomeScope 2 gives 224 Mb, below both, as expected where high-copy k-mers fall outside the fitted model. For *P. mocquerysi* the discrepancy is in the opposite direction, GenomeScope 2 returning 250.4 Mb against a 187.1 Mb assembly, a deficit that downward bias cannot explain. Comparing k-mer copy number against assemblies of the same reads that recover more sequence placed that deficit in repetitive sequence (Methods; Table S1). The deposited assembly collapses repeat arrays rather than omitting them, so its repeat content is approximately 36% against the 15.7% recovered, and repeat percentages for this species should be read as lower bounds.

### Gene Annotation

The GAGA annotation pipeline identified 10,860 representative protein-coding genes (21,665 total isoforms) in *P. mocquerysi* and 12,354 representative genes (25,317 isoforms) in *S. oregonense*. These counts are within the expected range for ant genomes, which typically contain 10,000–15,000 protein-coding genes (Vizueta et al., 2025). Compleasm recovered 98.2% complete BUSCOs for *P. mocquerysi* and 97.3% for *S. oregonense*, with fragmented fractions below 1% in both (Table 1), so the difference in contiguity does not translate into fragmented gene models.

### Repeat Content and Transposable Element Landscape

Repeat content differed dramatically between the two species: 15.7% in *P. mocquerysi* (29.1 Mb masked) versus 54.3% in *S. oregonense* (141.5 Mb masked; Table 2; Figure 3). The repeat landscape of *S. oregonense* is dominated by LTR retrotransposons (8.3% of the genome), DNA transposons (11.3%), and a large fraction of unclassified repeats (31.3%). In *P. mocquerysi*, unclassified repeats (11.8%) constitute the majority of the repeat fraction, with DNA transposons (1.8%) and LTR elements (1.1%) each contributing relatively little. This high amount of unclassified repeats represents the need for more accurate repeat classification in non-model organisms.

**Figure 3.**
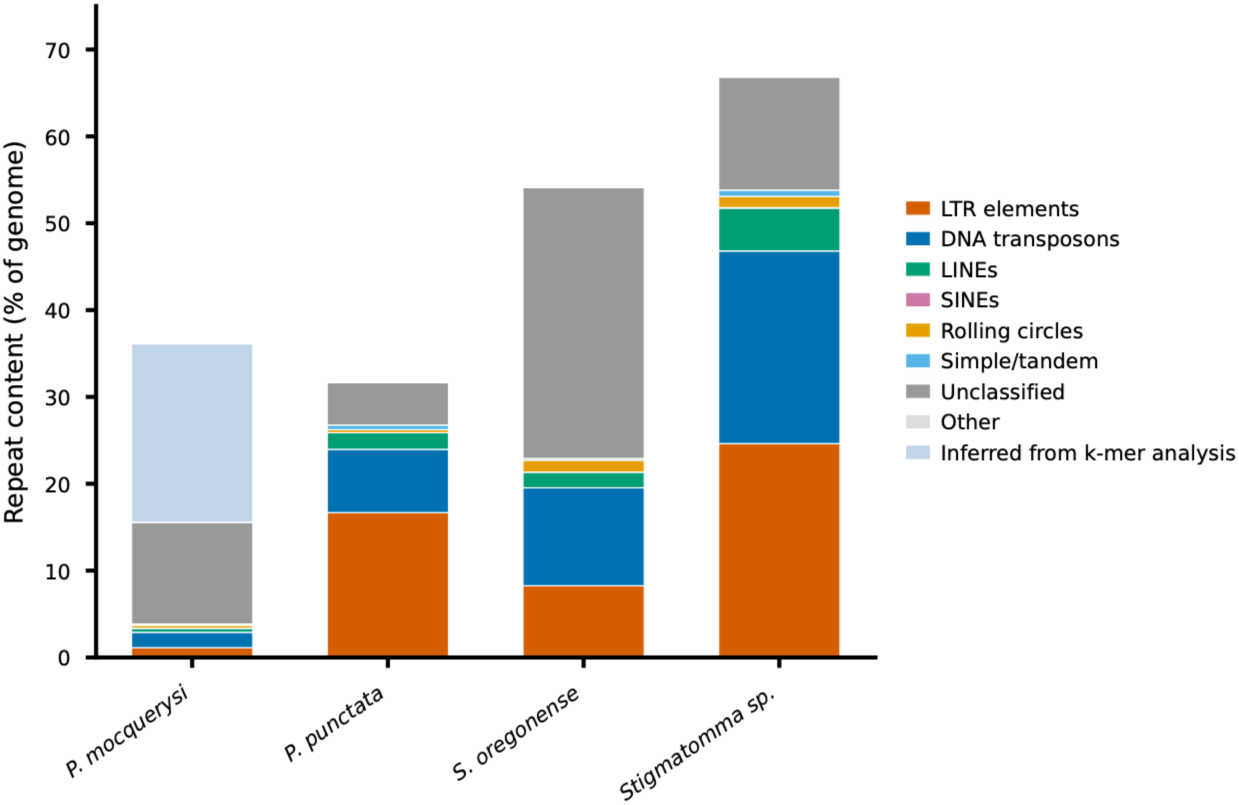
Repeat landscape showing the proportional contribution of major transposable element classes to each genome. LTR retrotransposons and DNA transposons dominate in Amblyoponinae, while unclassified repeats are the largest category in *P. mocquerysi*. The additional *P. mocquerysi* fraction is repeat content inferred from k-mer analysis of the reads, which the deposited assembly does not recover.

**Table 2.** Comparative genome statistics across four poneroid ant species. Repeat content was quantified from RepeatMasker annotations. *P. punctata* (GAGA-0090) and *Stigmatomma* sp. (GAGA-0391) assemblies and annotations are from the GAGA repository (Vizueta et al., 2025). Repeat percentages for P. mocquerysi are lower bounds; see Results.

|  | *P. mocquerysi* | *P. punctata* | *S. oregonense* | *Stigmatomma sp.* |
| --- | --- | --- | --- | --- |
| **GAGA ID** | GAGA-0601 | GAGA-0090 | GAGA-0602 | GAGA-0391 |
| **Subfamily** | Ponerinae | Ponerinae | Amblyoponinae | Amblyoponinae |
| **Genome size (Mb)** | 187.1 | 195.9 | 261.3 | 285.9 |
| **Scaffolds** | 307 | 285 | 1,189 | 587 |
| **N50 (Mb)** | 5.26 | 4.08 | 0.84 | 5.42 |
| **GC (%)** | 48.0 | 43.7 | 44.8 | 43.7 |
| **Genes (repr.)** | 10,860 | 11,340 | 12,354 | 13,182 |
| **Total repeat (%)** | 15.7 | 31.6 | 54.3 | 66.9 |
| **LTR elements (%)** | 1.1 | 16.7 | 8.3 | 24.7 |
| **DNA transposons (%)** | 1.8 | 7.3 | 11.3 | 22.3 |
| **LINEs (%)** | 0.5 | 1.9 | 1.8 | 5.0 |
| **Unclassified (%)** | 11.8 | 4.9 | 31.3 | 13.1 |
| **Rolling circles (%)** | 0.3 | 0.4 | 1.4 | 1.3 |

The unclassified fraction has three components. Of the 31.3%, 18.0% of the genome matched known transposable element families but could not be assigned to a superfamily with confidence, 7.6% comprised 565 de novo RepeatModeler families with no database homology, and 5.7% was tandem and simple repeats. The largest component is therefore diverged but recognisable element content, chiefly resembling hAT, Mariner/Tc1 and Polinton/Maverick families from other hymenopterans, left unclassified because Amblyoponinae is distant from the taxa with well-curated repeat libraries.

Profiling the same assemblies with longdust, an alignment-free low-complexity detector, identified 12.3 Mb of low-complexity sequence in *S. oregonense*, 4.69% of the genome and roughly double the other three species, driven by 205 arrays longer than 10 kb against 48 to 58 elsewhere (Table S2). The 20 most satellite-dense scaffolds contain no annotated genes. Low-complexity sequence thus accounts for only about 4.7 of the 31.3 percentage points, so the unclassified fraction consists predominantly of diverged transposable elements.

Comparison with the two additional poneroid species reveals a strong positive correlation between genome size and repeat content across the clade (Table 2; Figure 2). The two *Platythyrea* species have comparable repeat content once the collapse in the *P. mocquerysi* assembly is accounted for (approximately 36% against 31.6%; see Results and Methods). Amblyoponinae genomes (*S. oregonense*: 262 Mb, 54.3%; *Stigmatomma* sp.: 286 Mb, 66.9%) are substantially larger and more repeat-rich than Ponerinae genomes (*P. mocquerysi*: 187 Mb, 15.7%; *P. punctata*: 196 Mb, 31.6%). LTR/Gypsy elements are the single most abundant repeat family in all species with substantial repeat content, accounting for up to 16.5% of the *Stigmatomma* sp. genome (Figure 4).

**Figure 4.**
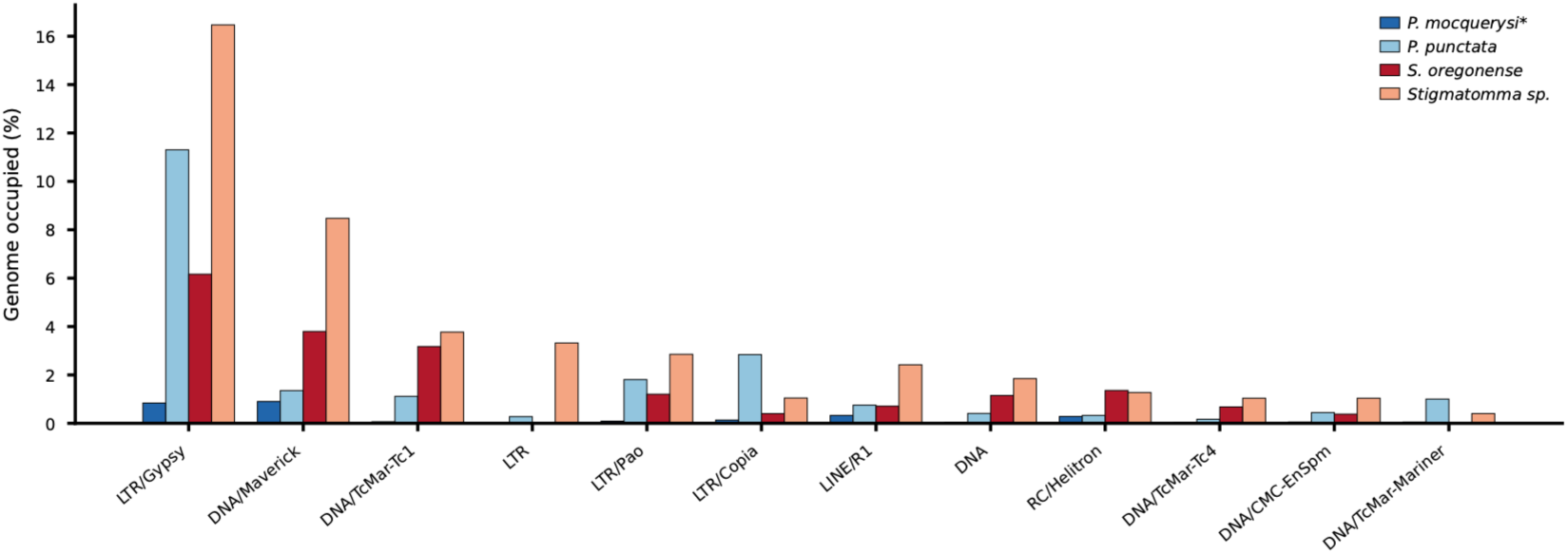
Top repeat families across species, showing the proportion of each genome occupied by the 12 most abundant specific repeat families. LTR/Gypsy is the dominant family in all species except *P. mocquerysi*. The asterisk marks *P. mocquerysi*, whose assembly collapses repeat arrays, so its values are lower bounds.

### Gene Family Variation

Comparison of BITACORA-annotated gene families across the four poneroid species reveals both conserved and variable repertoires (Table 3; Supplemental File S1). Several gene families are highly conserved: cuticle proteins (40–41 genes), vitellogenin (3 genes), DNMT methyltransferases (4 genes), serpins (7 genes), and GSTs (10–11 genes) are nearly invariant across all four species, suggesting strong functional constraint.

**Table 3.** Selected gene family counts across four poneroid ant species. Gene families were annotated using BITACORA v1.3 except for ORs and GRs, which were annotated by HAPpy-ABCENTH v1.0. Counts have not been manually curated. Full gene family data are provided in Supplemental File S1.

| **Gene family** | *P. mocquerysi* | *P. punctata* | *S. oregonense* | *Stigmatomma sp.* |
| --- | --- | --- | --- | --- |
| **Odorant receptors (OR)\*** | 342 | 408 | 382 | 396 |
| **OR fragments (pseudo)\*** | 12 | 0 | 43 | 0 |
| **Gustatory receptors (GR)\*** | 29 | 23 | 19 | 22 |
| **GR pseudogenes\*** | 0 | 1 | 2 | 1 |
| **Ionotropic receptors (IR)** | 34 | 27 | 35 | 27 |
| **Odorant binding proteins** | 10 | 16 | 15 | 19 |
| **Chemosensory proteins** | 11 | 11 | 13 | 21 |
| **Cytochrome P450s** | 69 | 84 | 82 | 82 |
| **Carboxylesterases** | 20 | 20 | 41 | 36 |
| **GSTs** | 10 | 10 | 10 | 11 |
| **cSerine proteases** | 7 | 7 | 16 | 15 |
| **Toll receptors** | 6 | 5 | 8 | 6 |
| **Serpins** | 7 | 7 | 7 | 7 |
| **Vitellogenin** | 3 | 3 | 3 | 3 |
| **Major royal jelly proteins** | 12 | 11 | 13 | 14 |
| **JH esterases** | 2 | 2 | 5 | 10 |
| **Fatty acid synthases** | 2 | 12 | 16 | 11 |
| **Acyl-CoA desaturases** | 7 | 9 | 20 | 21 |
| **Cuticle proteins** | 41 | 41 | 40 | 41 |
| **DNMT** | 4 | 4 | 4 | 4 |
| **Hexamerins/LSPs†** | 3 | 3 | 3 | 3 |

\*OR/GR annotated by HAPpy-ABCENTH v1.0; all other families by BITACORA v1.3. †Hexamerins identified by phylogenetic analysis (not in BITACORA pipeline; see text). For ORs and GRs, intact denotes models lacking the fragmentary identifier and includes some scored Partial or Pseudogene; that marker was not applied in the *P. punctata* and *Stigmatomma sp.* annotation runs, so fragment counts are not comparable across all four species.

In contrast, multiple gene families show subfamily-level expansions in Amblyoponinae relative to Ponerinae. Carboxylesterases are expanded from 20 genes in both Ponerinae species to 36–41 in Amblyoponinae. Clip-domain serine proteases (cSerine proteases), which play roles in innate immunity and melanisation, doubled from 7 in Ponerinae to 15–16 in Amblyoponinae. Lipid metabolism genes show particularly striking expansions: acyl-CoA desaturases expanded from 7–9 (Ponerinae) to 20–21 (Amblyoponinae), and fatty acid synthases from 2–12 to 11–16. Juvenile hormone (JH) esterases, which regulate JH titres and thus affect caste determination and reproduction, expanded from 2 (Ponerinae) to 5–10 (Amblyoponinae).

Cytochrome P450 detoxification genes are moderately expanded in *P. punctata* (84) relative to the other three species (69–82).

Hexamerins (also called larval serum proteins, LSPs) are haemolymph storage proteins derived from the hemocyanin superfamily and were not included in the standard BITACORA annotation pipeline. We identified three primary hexamerin/LSP orthologs in each of the four poneroid species, what we term hyLSP1, hyLSP2, and hyLSP3. These three genes are arranged as a syntenic tandem cluster conserved across all four species (*P. mocquerysi*: Scaffold13; *P. punctata*: Scaffold28; *S. oregonense*: Scaffold235; *Stigmatomma* sp.: Scaffold34). The hyLSP1 and hyLSP3 genes encode arylphorin-type hexamerins with high aromatic amino acid content (Phe+Tyr = 17–20% of residues), while hyLSP2 has moderate Phe+Tyr (∼12–13%) and elevated methionine content (6.6% in *P. mocquerysi*), consistent with a methionine-rich storage hexamerin. Hexamerins are of particular interest in these species given the central roles of haemolymph and larval secretions in poneroid ant nutrition and social regulation.

To establish that these expansions are not artefacts of the lower contiguity of the *S. oregonense* assembly we took a four-pronged approach (Table S3). Genes in the expanded families are no more likely than expected to lie near scaffold ends (4.9% and 5.0% against a null of 4.5%; exact binomial P = 0.56 and 0.61), although with 41 and 20 genes these tests would detect only gross enrichment. All 41 carboxylesterase and 20 acyl-CoA desaturase genes carry intact reading frames with canonical start and stop codons. Complete odorant receptors in *S. oregonense* have a median predicted length of 397 amino acids (aa), within the 392 to 398 aa range across the four species, so fragmentation has not systematically truncated them. The independently assembled *Stigmatomma* sp. genome shows matching family sizes (36 and 21).

### Chemosensory Gene Repertoires

The chemosensory gene repertoires of the four poneroid species are broadly comparable in size (Table 3). Odorant receptors (ORs), annotated by HAPpy-ABCENTH v1.0, ranged from 342 intact genes in *P. mocquerysi* to 408 in *P. punctata*, with the two Amblyoponinae species intermediate (382 in *S. oregonense*; 396 in *Stigmatomma* sp.). These counts are within the range reported for other ant genomes in the GAGA consortium (Vizueta et al., 2025). *S. oregonense* had a notably higher number of OR pseudogene fragments (43) compared to the other species (0–12), which likely reflects assembly fragmentation of tandem OR arrays given its lower contiguity (N50 = 0.84 Mb). The intact models show no sign of truncation. Complete ORs have median predicted lengths of 392, 393, 397 and 398 aa across the four species, and of the 317 scored Complete in *S. oregonense* all retain an intact reading frame (Table S3).

Gustatory receptors (GRs) were more variable, ranging from 19 intact genes in *S. oregonense* to 29 in *P. mocquerysi*. Ionotropic receptors (IRs) were relatively conserved across species (27–35 genes), as were odorant-binding proteins (OBPs; 10–19) and chemosensory proteins (CSPs; 11– 21), with a trend toward higher counts in Amblyoponinae. The overall conservation of chemosensory repertoire sizes across these four species suggests that poneroid ants maintain large OR complements regardless of their divergent life histories and ecological niches.

### Limitations and Future Directions

Retrieving high-quality genome assemblies from limited individuals as the tiny *S. oregonense* ants remains a challenge. For instance, species with limited biomass available targeted by the GAGA project were unable to provide long-read sequencing data, preventing the generation of contiguous assemblies. Therefore, further sampling across the ant tree of life, and the development of ultra-low input sequencing methods, will allow the generation of genome data to cover additional representation of the diverse life history and social traits across ants.

The gene family annotations presented here were generated as part of a high-throughput pipeline applied across hundreds of ant genomes (Vizueta et al., 2025) and have not been manually curated for these species. While the automated annotations provide a useful first approximation, further curation or validation (as used in Pellen et al. 2026) would refine the gene counts reported here. The 43 OR pseudogene fragments in *S. oregonense* likely reflect its lower assembly contiguity rather than true pseudogenisation, though the intact models themselves show no evidence of truncation. Future work should include: (i) further curation and validation of gene annotations, e.g. supporting chemosensory gene family members with antennal transcriptome data; (ii) the annotation and classification of uncharacterised repeats; (iii) chromosome-level scaffolding of the *S. oregonense* assembly, which would improve resolution of tandem gene arrays; and (iv) functional genomic studies leveraging these assemblies to investigate the molecular basis of larval secretion in *P. mocquerysi* and haemolymph feeding in *S. oregonense*.

## Data Availability

The genome assemblies, raw sequencing reads, and annotations have been deposited at NCBI under BioProject PRJNA1434519. The *Platythyrea mocquerysi* genome assembly (BioSample SAMN56973653) is available under WGS accession JBWYQZ000000000, and the *Stigmatomma oregonense* genome assembly (BioSample SAMN56973654) is available under WGS accession JBWYQY000000000. Genome annotations (GFF3, CDS and protein FASTA) and RepeatMasker output are archived at Zenodo under DOI 10.5281/zenodo.19595400. Detailed gene family counts are provided in Supplemental File S1. Raw genome sequencing reads are deposited under BioProject PRJNA1513708. Custom analysis code is available at https://github.com/Social-Fluids-Lab/Poneroid_Genomes and archived at Zenodo under DOI 10.5281/zenodo.22705568; the GAGA annotation pipeline is at https://github.com/schraderL/GAGA. The voucher specimens (CASENT0872487, CASENT0872488) are deposited at the California Academy of Sciences.

## Acknowledgements

We thank Pamela Nicholson and the Next Generation Sequencing Platform at the University of Bern for PacBio sequencing, the Madagascar Biodiversity Center team for support in the field, and the GAGA consortium for providing the annotation pipeline and comparative data. We thank Alexander Wild and François Brassard for the photographs in Figure 1. During the preparation of this work, Claude (Anthropic, models 4.7-5) was used to generate and improve readability of the code used to produce the figures and to improve readability of the manuscript. After using this tool, the authors reviewed and edited the code and content and take full responsibility for the content of the publication.

## Supplementary Material

Supplemental File S1. Gene family annotations for the four poneroid species (Gene_Family_Details.xlsx). Per-species sheets list the individual gene models recovered by BITACORA and HAPpy-ABCENTH, with summary, cross-species, chemosensory and hexamerin sheets.

### Tables

**Table S1.** Assemblies and assembly-free genome size estimates for *P. mocquerysi*. The Wtdbg2 assembly is the one deposited; the Hifiasm haplotypes were generated during assembler selection. BUSCO figures are Compleasm in genome mode against hymenoptera_odb10, the dataset used when these assemblies were compared; the protein-mode values in Table 1 are against hymenoptera_odb12. Longest LC array is the longest low-complexity array reported by longdust v1.4 (-k 8 -w 20000).

| Assembly or estimate | Span (Mb) | Contigs | N50 (Mb) | GC (%) | Duplicated BUSCO | Longest LC array (kb) |
| --- | --- | --- | --- | --- | --- | --- |
| Wtdbg2 (deposited) | 187.1 | 307 | 5.26 | 48.04 | 0.33% | 52 |
| Hifiasm haplotype 1 | 297.0 | 979 | 0.90 | 46.65 | 9.93% | 260 |
| Hifiasm haplotype 2 | 254.2 | 995 | 0.61 | 46.85 | 5.54% | 214 |
| GenomeScope 2 estimate | 250.4 | - | - | - | - | - |

**Table S2.** Low-complexity sequence content across the four assemblies (longdust v1.4, -k 8 -w 20000).

| Species | Assembly (Mb) | Low-complexity (Mb) | % genome | Arrays >10 kb | Max (kb) |
| --- | --- | --- | --- | --- | --- |
| *S. oregonense* | 261.3 | 12.3 | 4.69 | 205 | 347 |
| *Stigmatomma* sp. | 285.9 | 5.7 | 2.01 | 58 | 92 |
| *P. mocquerysi* | 187.1 | 4.6 | 2.44 | 50 | 52 |
| *P. punctata* | 195.9 | 3.1 | 1.59 | 48 | 50 |

**Table S3.** Boundary proximity and coding sequence integrity of expanded gene families in *S. oregonense*. Null expectation for boundary proximity is 4.5%. Odorant receptors are given for all models and for those scored Complete.

| **Gene family** | **n** | **Near boundary** | **ATG start** | **Canonical stop** | **Frame intact** |
| --- | --- | --- | --- | --- | --- |
| CCE | 41 | 2 (4.9%) | 41 (100%) | 41 (100%) | 41 (100%) |
| Desat | 20 | 1 (5.0%) | 20 (100%) | 20 (100%) | 20 (100%) |
| OR, all models | 425 | 11 (2.6%) | 288 (67.8%) | 377 (88.7%) | 407 (95.8%) |
| OR, complete only | 317 | 5 (1.6%) | 288 (90.9%) | 316 (99.7%) | 317 (100%) |

